# A new species of leaf fish, *Nandus banshlaii* (Perciformes: Nandidae) from West Bengal, India

**DOI:** 10.1101/2020.08.02.232751

**Authors:** R. Kapuri, A. K. Sinha, P. de, R. Roy, S. Bhakat

**Affiliations:** Department of Zoology, Rampurhat College, Rampurhat 731224, West Bengal, India

**Keywords:** Banshlai River, Black spot, Gangetic delta, Head length, Heatmap, PCA

## Abstract

*Nandus banshlaii*, sp. nov. is described from the Banshlai River of West Bengal. This species is distinguished from all its congeners in having a golden brown body in live and a combination of characters like longest head and snout length (44.28% SL and 35.58% HL respectively) and from its two Indian congeners in containing largest eye diameter (22.22% HL), longest pre-dorsal length (47.16% SL), shortest pectoral fin length (14.60% SL) and smallest dorsal fin base length (45.53% SL).Like all three congeners of Ganga-Brahmaputra-Meghna river basin of Gangetic delta it possess a dark spot on caudal peduncle.

To differentiate the present species from other two Indian congeners of *Nandus*, *N. nandus* and *N. andrewi*, PCA and heatmap is performed and a key of all three species is also provided.

## Introduction

Genus *Nandus* Valenciennes in Cuvier and Valenciennes, 1831, commonly known as leaf fish is characterized by oblong compressed body, large protrusible mouth with projecting lower jaw, triangular operculum with a single prominent spine, dorsal fin with 12 – 14 spines and 11 – 13 soft rays, anal fin with three spines and seven to nine soft rays, slightly rounded caudal fin, ctenoid scale, interrupted lateral line and cryptic colouration. The fresh water perciforms are distributed throughout South and South-east Asia in both lentic and lotic hahitats.

Seven nominal species are currently recognized in their natural distributional ranges, namely *Nandus nandus* (Hamilton, 1822) from Indian subcontinent and Myanmar, *N. andrewi* (Ng and Jaafar, 2008) from the Ichamati river drainage in North-east India, *N. nebulosus* (Gray, 1835) from the Malay Pennisular, Sumatra and Borneo, *N. oxyrhynchus* (Ng, Vidthayanan and Ng, 1996) from Mekong river drainage in central Thailand, *N. prolixus* (Chakraborty, Oldfield and Ng, 2006) from Borneo, Malaysia, *N. mercatus* from Musi river drainage in Southern Sumatra (Ng, 2008) and *N. meni* from Noakhali, Bangladesh (Hossain and Sarker, 2013). Two other species of *Nandus, N. marmosratus* Cuvier and Valenciennes (1831) and *N. borneensis* (Steindachner, 1901) are conspecific of *N. nandus* (Day, 1878) and *N. nebulosus* (Weber and de Beaufort, 1922) respectively.

The leaf fish mainly inhabits in streams, rivers, pools, ponds, ditches, inundated fields, reservoirs (Hamilton, 1822; Talwar and Jhingran, 1991 and Rainboth, 1996). Once it was very common in West Bengal (India) and in summer months; plenty of fish is collected from dried-up beds of reservoirs, tanks, beels, bheries etc. It had great demand in the market due to its taste. But now it is seldom available in the market as its natural population has declined to an alarming extent in this region. As a result this fish has been categorized as critically endangered by Das and De, 2002. In Bangladesh this fish is treated as vulnerable (IUCN Bangladesh, 2000), although it is placed as least concern globally (IUCN 2014).

Since 2014, the corresponding author is collecting different species of fish from rivers, ponds, beels, canals, reservoirs etc. either directly from the natural habitat or from fish markets of Birbhum district, but he never found any specimen of *Nandus* sp. so far. Accidentally, the first author recovered 20 specimens from Banshlai River on 27. X. 2019. The river was flooded due to incessant rain (on and from 23. X. 2019 to 26. X. 2019) and plenty of different species of fishes were available. It is also interesting to note that the local people including the fishermen in this area are unable to identify the very fish. The findings of above mentioned authors reflect that *Nandus* sp. population in Birbhum district is rare.

The present paper describes this newly recovered *Nandus* sp. from Banshlai River as a new species, *Nandus banshlaii*.

## Materials and Methods

The specimens collected from Banshlai River were preserved in 10% formalin. Before preservation colour pattern of the live fish was studied carefully. All the specimens of this species were brought to the Department of Zoology, Rampurhat College for taxonomic identification. Measurements were made point to point for each specimen with dial caliper to the nearest 0.1mm accuracy. Counts and measurements were made on the left side of the specimens. Fin ray and scale counts and other details of specific characters were made with the help of stereo zoom microscope.

Measurements of different body parts including head length (HL) are given as percentage of standard length (SL) and the subunits of head are presented as percentage of the head length. All the measurements and counts are followed by Hubbs and Lagler (2004)

To separate different Indian species of *Nandus*, Principal Component Analysis (PCA) and heat map are performed using the data presented in Table 1.

**Table 1.** Biometric data of *N. banshlaii* (n= 20).

| Parameters | Holotype | Range | Mean± SD |
| --- | --- | --- | --- |
| Total length (TL in mm) | 68.00 | 45.51-69.50 | 57.70 |
| Standard length (SL in mm ) | 54.20 | 38.10-58.52 | 47.52 |
| % SL |  |  |  |
| Pre- dorsal length (PDL)* | 46.12 | 43.63-50.00 | 47.16± 1.27 |
| Pre-pectoral length (PPL)* | 40.22 | 39.46-50.00 | 43.89± 2.75 |
| Pre ventral length (PVL)* | 42.43 | 39.00-49.87 | 43.98± 1.33 |
| Pre anal length (PAL)* | 76.57 | 71.11-79.80 | 74.92± 1.14 |
| Dorsal fin length (DFL) | 16.24 | 13.63- 20.00 | 16.69±1.80 |
| Pectoral fin length (PFL)* | 14.76 | 12.98- 19.95 | 14.60± 1.91 |
| Ventral fin length (VFL)* | 16.79 | 14.56- 21.11 | 17.44± 1.58 |
| Anal fin length (AFL) | 15.68 | 12.72- 19.44 | 15.85± 1.33 |
| Anal fin spine length (AFSL)* | 9.01 | 7.76- 11.11 | 9.11± 1.31 |
| Dorsal fin base length (DFBL)* | 46.86 | 41.41- 50.00 | 45.53± 2.94 |
| Pectoral fin base length(PFBL) | 7.38 | 5.98- 7.95 | 6.87± 0.57 |
| Ventral fin base length (VFBL) | 3.32 | 3.44- 5.55 | 4.28± 0.62 |
| Anal fin base length (AFBL) | 12.73 | 8.49- 12.56 | 10.97± 1.15 |
| Pectoral fin to Ventral fin distance | 3.32 | 2.83- 4.44 | 3.14± 1.05 |
| Ventral fin to Anal fin distance | 34.02 | 27.27- 35.52 | 31.11± 2.08 |
| Vent to Anal fin distance | 6.08 | 3.96- 6.83 | 5.26± 0.87 |
| Head length (HL)* | 44.38 | 38.61- 46.13 | 44.28± 1.38 |
| Head Width (HW) | 11.07 | 9.81- 13.69 | 11.97± 0.89 |
| Head depth ( HD) | 27.31 | 25.31- 29.16 | 26.72± 0.88 |
| Caudal peduncle length (CPL)* | 11.99 | 11.84- 16.36 | 13.92± 0.81 |
| Caudal peduncle depth (CPD)* | 12.32 | 10.52- 13.88 | 11.58± 0.57 |
| Body depth at anus (BD)* | 28.04 | 24.93- 32.00 | 28.99± 1.44 |
| Body depth at dorsal fin | 34.69 | 30.26- 37.86 | 34.75± 0.97 |
| Caudal fin length (CFL)* | 23.24 | 17.27- 25.00 | 21.32± 1.86 |
| % HL |  |  |  |
| Eye diameter (ED)* | 20.17 | 16.21- 26.66 | 22.22± 1.87 |
| Inter orbital distance (IOD)* | 24.45 | 21.62- 29.62 | 26.61± 2.05 |
| Inter narial distance (IND) | 13.92 | 13.51- 17.64 | 15.32± 1.79 |
| Gape | 29.25 | 28.03- 34.98 | 30.48± 1.27 |
| Snout (SN)* | 33.68 | 32.35- 44.11 | 35.58± 1.67 |

\* Used for PCA and heatmap

PCA is a statistical process where a high dimensional multivariate dataset is linearly transformed into a set of uncorrelated low dimensional linear variables called principal component. In PCA, first few (usually first two PC1 and PC2) principal component often explain large amount of variations.

Heatmap is a graphical presentation of data-matrix where individual values contained in a block. The data contained within a block is based on the relationship between two variables connecting the row and column. Here categorical data is colour coded and numerical data is presented in colour scale that blends from one colour to another in order to represent the differences in high and low value. The dendrogram along both sides of heatmap show how the variables (species) and the rows (characters) are independently clustered. Both PCA and heatmap analysis are performed using on line graphical user interface Clust Vis (Jollife, 2002; Metsalu and Vilo, 2015).

To describe the shape of snout, angle of snout is measured by extending the line from lower jaw and head in the photographic impression of seven species of *Nandus* (five species from Ng and Jaafar, 2008 and *N. mercatus* from Ng, 2008). The angle ≤ 60° indicates pointed snout and > 60° for blunt snout. Moreover, a formula is presented to measure the shape of the snout

> Shape of snout = Length of pre-orbital vertical line (a) / Length of snout (b).
>
> The value ≤ 1.4 indicates pointed and higher value indicates blunt.

## Results

***Nandus banshlaii* sp. nov.** (Plate 1).

### Type material

Holotype: Hamilton Museum of Fresh Water Fishes (HMFWF – PN86/2019), Rampurhat College, Rampurhat-731224, District Birbhum, W. B. India. 54.20 mm SL; collected from Banshlai River at Palsa, (24°27’49” N 87°51’00” E) Rampurhat Subdivision, Birbhum district, West Bengal, India, by R. Kapuri, 27 October, 2019.

Paratype: 19 ex., 35.5 – 57.0 mm SL. All other details are same as holotype.

### Diagnosis

*Nandus banshlaii* is distinguished from all its congeners by longest head length (44.28% SL vs. 39.4% SL in *N. prolixus* to 43.1% SL in *N. oxyrhynchus*) including Indian congeners (39.5% SL in *N. nandus* and 41.6% SL in *N. andrewi*). Body depth of *N. banshlaii* is almost same as *N. andrewi* (28.99% SL in present species and 29.1% SL in *N. andrewi*), but smaller in respect to in *N. nandus* (33.9% SL). Head width is slightly greater to *N. nandus* (11.97% SL in *N. banshlaii* and 10.19% SL in *N. nandus*), but smaller in respect to that of *N. andrewi* (14.5% SL). Eye is distinctly larger from its Indian congeners like *N. nandus* and *N. andrewi* (eye diameter 22.22% HL in *N. banshlaii* whereas, in *N. nandus* and *N. andrewi* the values are 17.2% and 21.7% of HL respectively). Snout length is maxima in *N. banshlaii* compared to all other species (35.58% HL vs. 26.1% HL in *N. nebulosus* to 30.0% HL in *N. oxyrhynchus*). Pre-dorsal length in *N. banshlaii* is maxima compared to other two Indian species (47.16% SL vs. 42.20% SL in *N. nandus* and 41.70% SL in *N. andrewi*). Pre-pectoral length in *N. nandus* and *N. andrewi* is almost similar (39.80% SL) but in the present species it is longest (43.89% SL). Pectoral fin length in *N. banshlaii* is shortest (14.60% SL) compared to *N. nandus* (17.40% SL) and *N. andrewi* (17.00% SL). Among all three *Nandus* species of Gangetic River basin dorsal fin base length is minimum in the present species (45.53% SL vs. 48.50% Sl in *N. nandus* and 51.00% SL of *N. andrewi*.). Caudal peduncle depth in caudal peduncle length is maximum in *N. banshlaii* (1.20) compared to other two Indian congeners *N. nandus* (1.15) and *N. andrewi* (1.07).

Most distinguishing and also diagnostic character in this species of *Nandus* is the arrangement of lateral line. Two lateral lines (interrupted) upper and lower may overlap or not. The gap between these two lines is of three to three and half scales different. Moreover, the number of lateral line scales varies widely. It ranges from 36 to 50. But in other two Indian species the number of lateral line scales varies from 42 – 55 in *N. nandus* and 45 – 52 in *N. andrewi*.

### Description

The morphometric data for *N. banshlaii* is presented in Table I. Body is laterally compressed and moderately elongate. Dorsal profile of the body evenly slopes down but the ventral profile is almost straight from isthmus to anal fin origin. Head is also compressed and sharp. Snout is triangular and acute. Mouth is moderately large, protrusible and with deep cleft, superior, strongly oblique. Posterior end of maxilla extends beyond orbit. A thin soft Y-shaped cover is present on both the jaws. Eye is large, circular and present in the upper middle half of the head. Snout tip to isthmus is straight. Posterior edge of preopercle is finely serrated. Upper portion of opercle bears a small spine. In the dorsal portion of the head, there is a long rod like ridge in the middle extending from anterior end of nostril to the middle of head. There are two lateral lines, the anterior extending from the tip of the operculum to the middle of the anal fin base and the posterior from the middle of anal fin base to the caudal peduncle.

Dorsal fin inserted at vertical through middle of pectoral fin length. Origin and ending of soft dorsal and anal fin are almost parallel and tip of both fins almost touch caudal peduncle. Dorsal fin base is longest and covers nearly half of standard length. Ventral fin origin is on the anterior one third of pectoral fin length. Caudal peduncle depth in caudal peduncle length is 1.20.

Dorsal fin contains 12 – 13 spines, the edge of which is dented like saw but the edge of soft fin is roundish and contains 9 to 10 rays. Spiny part is longer than soft part. Pectoral fin, the smallest fin, 14.60% SL and is inserted in the lower one fourth of flank just behind operculum. It contains 14 – 16 rays and without spine. Ventral fin is dagger shaped (when closed) with small base (only 4.82% SL) and contains one spine with five soft rays. Anal fin with three spines, first one is the smallest and number of soft fin rays 6 to 7. Like dorsal fin, anal fin edge is also roundish. Caudal fin is 25.60% SL with 13 to 14 rays. A black round spot is on the middle of caudal peduncle surrounded by whitish circle followed by two blackish rhomboid structures.

Scales are ctenoid, imbricate and nearly uniform in size, present throughout the body including head except a few region viz. in the upper lip, maxillary area, gular region, mid-dorsal ridge of head and inter-orbital line. Number of lateral line scales varies from 36 to 50 of which anterior line bears 26 to 35 scales and posterior line bears 10 to 15 scales.

Numerous villiform teeth are present on jaws, palate and tongue. The tongue is free and with two distinct longitudinal ridges.

Fin rays count:

DF XII – XIII, 9 – 10; PF 14 – 16; VF I, 5; AF III, 6 – 7; CF 13 – 14; Branchiostegal rays 6.

Scale count:

Lateral line scale 36 – 50, Pre-dorsal scale 12 – 15, Circumpeduncular scale 25 – 28, above lateral line 5 – 6, below lateral line 13 – 15.

### Colouration

In live specimen, the colour of the fish is reddish from the tip of the snout to the base of the operculum except the posterior portion of the eye while it is reddish black in the dorsum but golden brown colouration is observed in the flank. Reddish lunar shape structure is present in the base of the pectoral fin and operculum. Round black spot is present on the caudal peduncle. Three irregular patches from the top to the bottom of the flank in the posterior one-third portion of the body and two irregular patches in the middle are seen both of which are below the anterior lateral line in the middle of the body.

In the preserved specimen (in 10% formalin solution), three distinct dark longitudinal patch of irregular shape running across the body from the dorsum to the ventrum. Ventral position of the body is lighter. Lower portion of mandible is whitish.

All the fins are with spotted bands except pectoral fin which is transparent. Both soft dorsal and anal fin contains three narrow lined dark bands along its length. In the ventral fin proximal two-third portion bears blackish band though distal one-third is hyaline. Caudal fin is with four blackish zigzag bands. Fin membranes of spiny rays are hyaline. Operculum is with whitish patch that extends posterior to orbit and below it. Caudal peduncle is with a black round spot encircled by whitish circular ring.

Etymology

The name banshlaii is latin referring to its collection site, Banshlai River.

### Distribution

Currently known from its type locality, Banshlai River, Birbhum district, West Bengal. The Banshlai River is the first tributary of Bhagirathi of Gangetic River basin. Banshlai River flows through Pakur district of Jharkhand and Birbhum and Murshidabad district of West Bengal before entering the Bhagirathi at north of Jangipur (Murshidabad district). The river covers a distance of 112 km of which only 17% is within West Bengal.

### Habitat

*N. banshlaii* was collected in slow, moderate running water consisting of various substrata such as mud particle, less amount of sand particles and algae and other herbs on the substratum in the dry season. A few deep small stagnant water bodies are present here and there in the river bed throughout the dry season (October to May) and sometimes used as seed bed for paddy culture (Fig. 2A and 2B). But in the rainy season when the fish was collected, a flush of rapid and turbid flowing water is observed. Other associated fish species were *Labeo calbasu, Puntius ticto, P. sarana*, *Osteobrama cotio*, *Amblypharyngodon mola* (Cyprinidae), *Arius arius* (Ariidae), *Mystus vittatus*, *M. cavasius* (Bagridae), *Wallago attu, Ompok pabo* (Siluridae), *Bagarius bagarius, Gagata cenia* (Sisoridae) etc.

**Fig.1.**
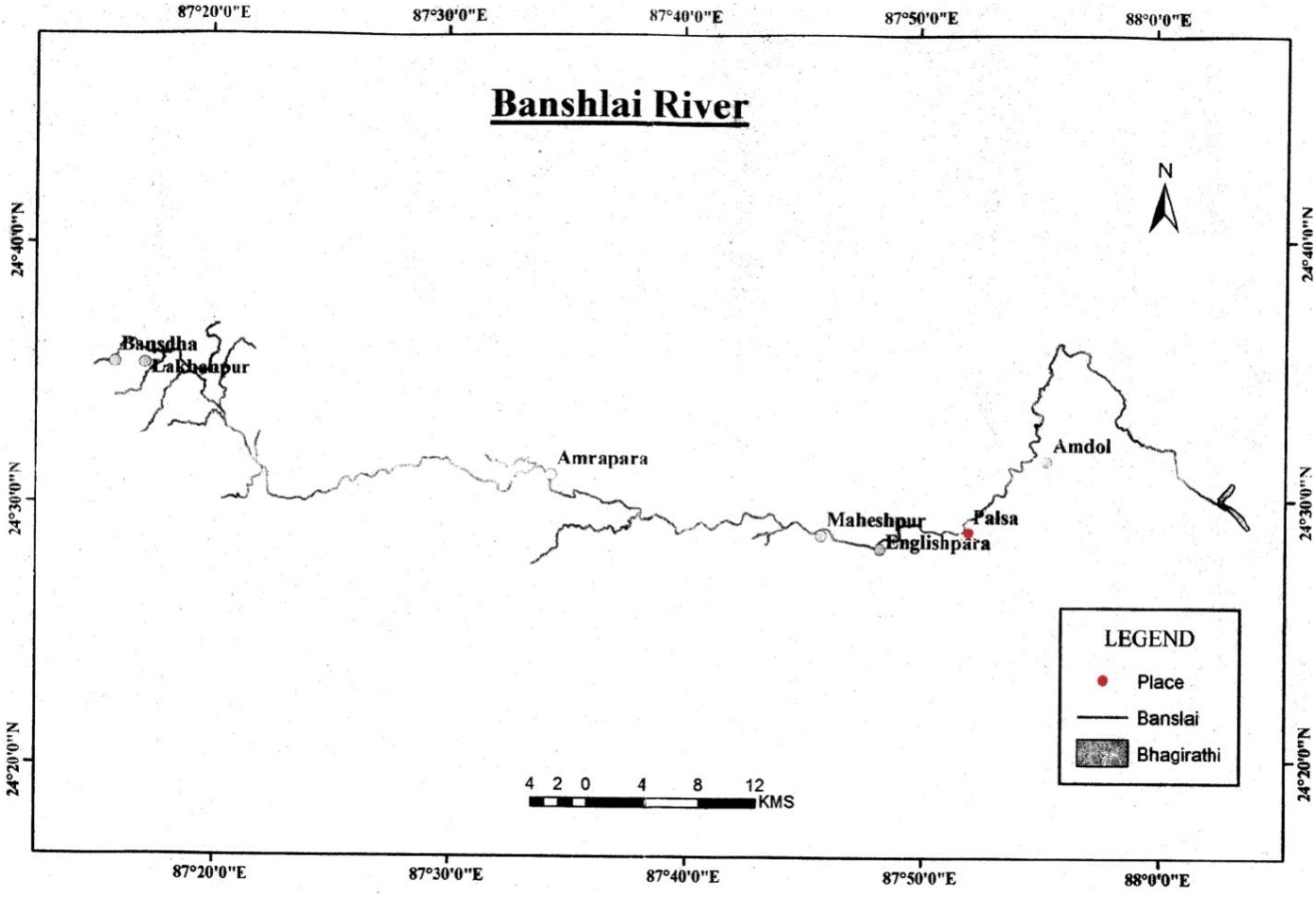
Sampling site at Palsa (marked as •) of Banshlai river.

**Fig. 2A:**
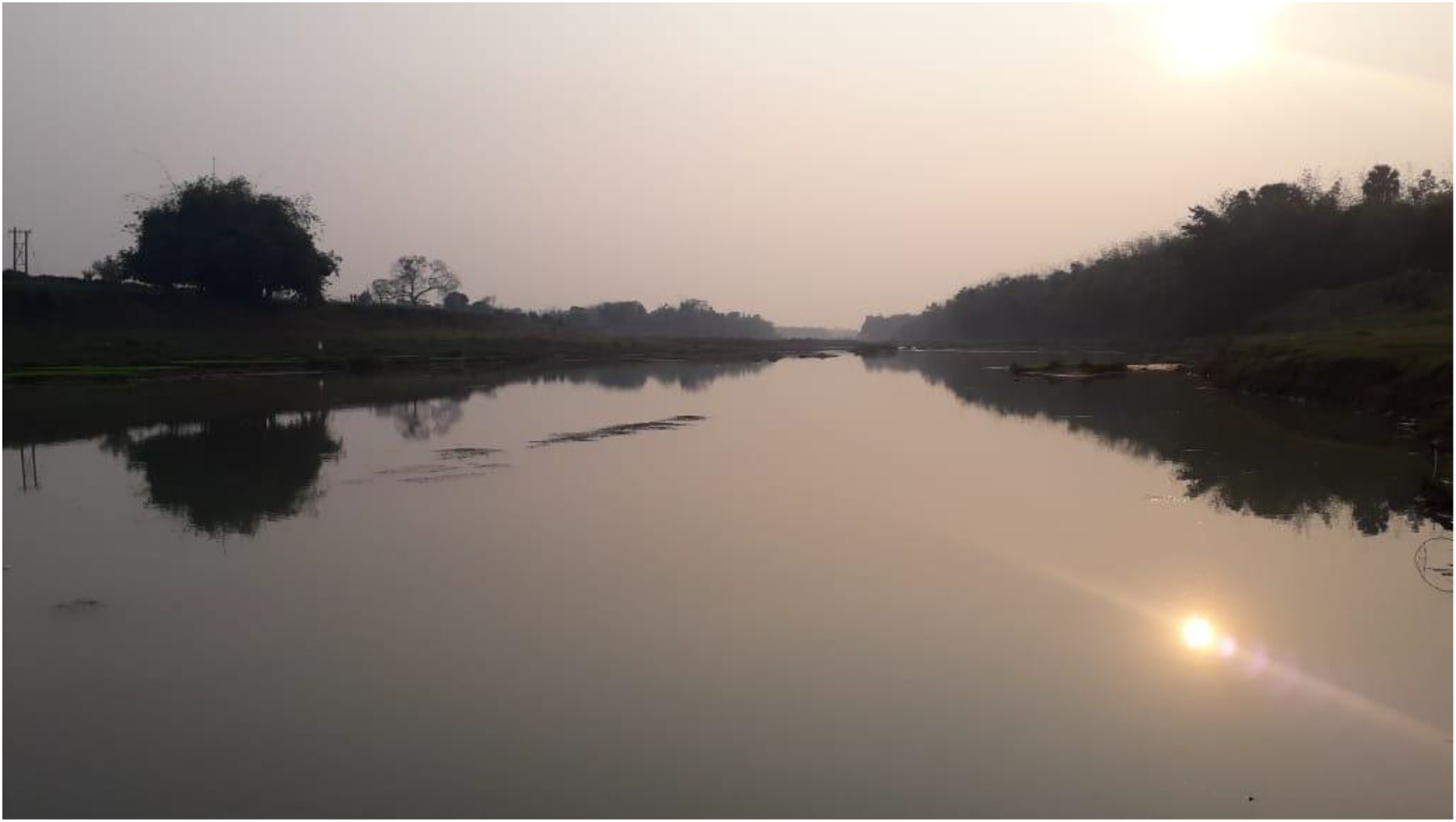
Banshlai River Bed

**Fig. 2b.**
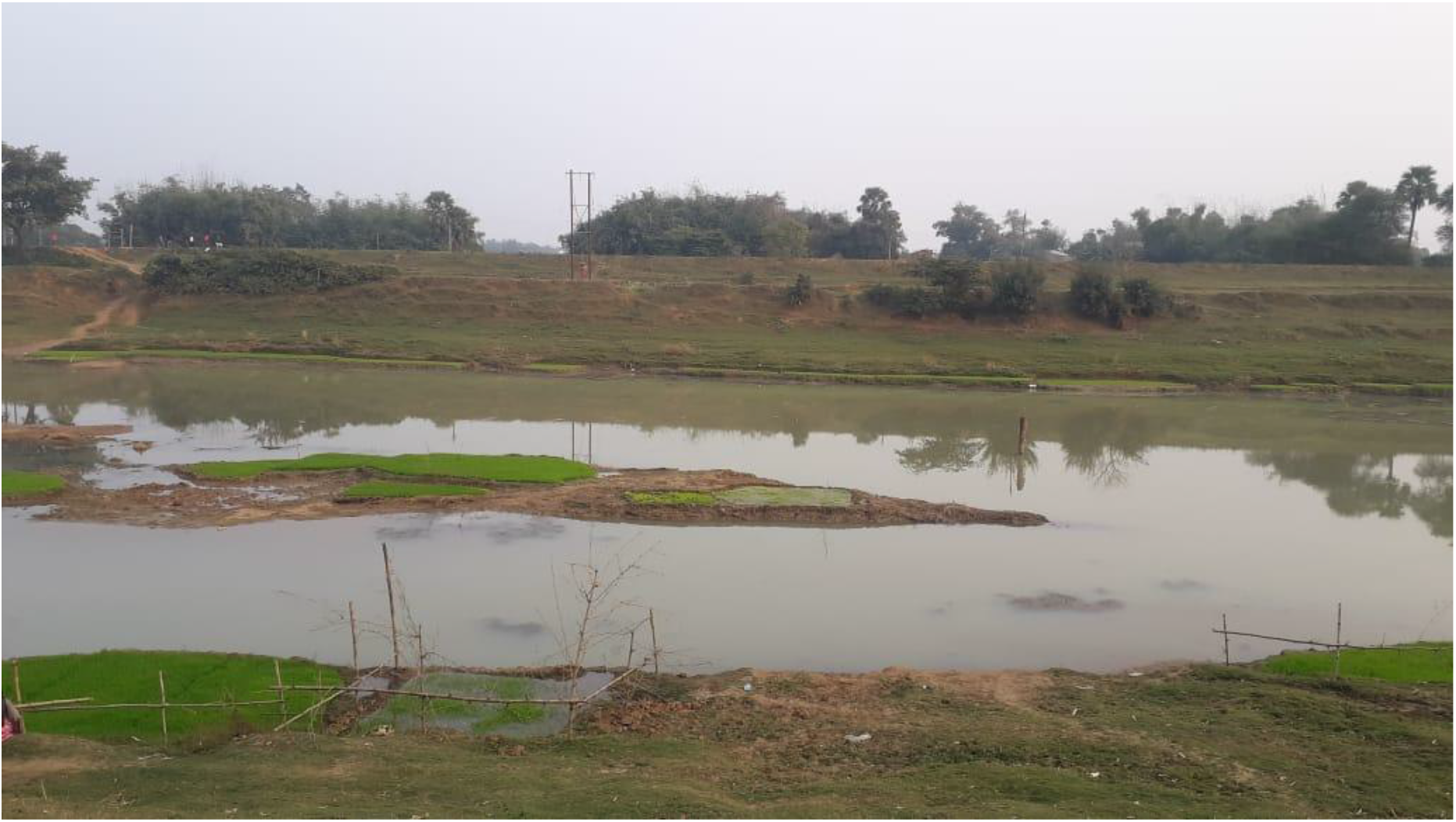
Banshlai River Bed (Strictly in dry season)

### Comparison

Snout of all Indian congeners viz. *N. nandus, N. andrewi* and *N. banshlaii* is pointed (angle of snout ≤ 60° and shape of snout ≤ 1.4, but in other non-Indian species, *N. nebulosus, N oxyrhynchus, N. prolixus* and *N. mercatus* the snout is comparatively blunt (Table 2A). Among all the eight species, head length of *N. banshlaii* is maximum (44.28% SL) whereas, in other species it ranges from 39.4% SL (*N. prolixus*) to 43.1% SL (*N*. *oxyrhynchus*), but body depth is lowest (28.99% SL) and nearest to *N. andrewi* (29.1% SL). *N. banshlaii* has longest snout (35.58% HL vs. 26.1% HL to *N. nebulosus* to 30.0% HL in *N. oxyrhynchus*) (Table 2A). Caudal peduncle depth in caudal peduncle length in three Indian species is > 1.00 but in other non Indian species it is ≤ 1 (0.76 – 0.86 in *N. mercatus*, 0.99 in *N. prolixus*, and 1.04 in *N. meni*).

**Table 2A.** Comparison of seven species of *Nandus sp*. (Hossain and Sarker, 2013) with *N. banshlaii*.

| Parameters | *N. banshlaii* * | *N. meni* | *N. nandus* * | *N. oxyrhynchus* | *N. andrewi* * | *N. mercatus* | *N. nebulosus* | *N. prolixus* |
| --- | --- | --- | --- | --- | --- | --- | --- | --- |
| SL (mm) | 57.00 | 111.90 | 127.70 | 54.60 | 125.00 | 70.00 | 83.80 | 83.40 |
| HL (%SL) | 44.28 | 40.40 | 33.90 | 43.10 | 41.60 | 39.70 | 41.30 | 39.40 |
| BD (%SL) | 28.99 | 30.30 | 33.90 | 43.80 | 29.10 | 43.00 | 43.70 | 39.00 |
| HW (%SL) | 11.97 | 12.24 | 10.19 | 18.20 | 14.50 | 20.00 | - | 18.20 |
| ED (%HL) | 22.22 | 18.90 | 17.20 | 26.00 | 21.70 | 23.00 | 33.10 | 25.40 |
| Snout (%HL) | 35.58 | 29.70 | 29.90 | 30.00 | 28.60 | 27.00 | 26.10 | 27.10 |
| Shape of Snout | 1.343 | - | 1.40 | 1.713 | 1.425 | 1.619 | 1.833 | 1.483 |
| Angle of Snout | 60° | - | 60° | 72° | 59° | 70° | 77° | 65° |

\*Indian species

Number of pre-dorsal scales in *N. banshlaii* and *N*. *meni* is minimum (only 12) compared to other five species. Dark spot on caudal peduncle is present only in all three Indian congeners along with *N. meni* (species from Bangladesh). Number of scales above lateral line is similar in all three Indian species of *Nandus* including the species of Thailand (*N. oxyrhynchus*), though that of below lateral line is variable. Among Indian species of *Nandus*, number of circumpeduncular scales is minimum in *N. andrewi* compared to other two species (21vs. 26 in both *N. nandus and N. banshlaii*) and maximum in a species from Bangladesh, *N. meni* (Table 2B).

**Table 2B.**
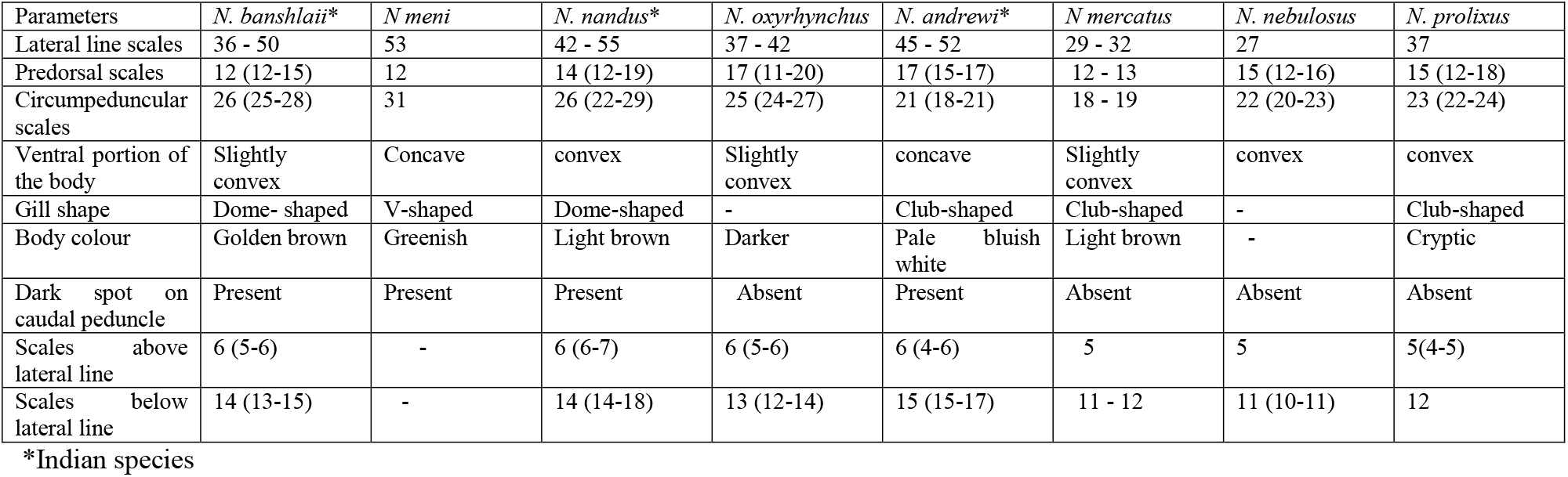
Comparison of seven species of *Nandus sp*. (Hossain and Sarker, 2013) with *N. banshlaii*.

Body colour of the present species is golden brown and cryptic but in *N. nandus* it is light brown whereas, in *N. andrewi* it is pale bluish white. Number of soft fin rays of dorsal fin in the present species is minima (9 – 10), whereas, it varies from 11 – 13 and 11 – 14 in *N. nandus* and *N. andrewi* respectively. Number of pectoral fin rays and ventral fin rays are almost equal in all three Indian congeners of *Nandus*. Number of caudal fin rays in all the eight species of *Nandus* is almost same. In case of other fins, the number of ray is variable. Though number of spines in dorsal fin in all the eight species of *Nandus* are variable (ranges from XII to XIV), while that of anal fin is constant (only III) (Table 3).

PCA provides a summarization of 17 morphological characters among three species of Indian congeners of *Nandus* (Fig. 3). Principal component (PC) 1 explains 74.8% variation among species and PC2 explains 25.2%. On the basis of PC1, all three species are equally and distantly related though *N. andrewi* and *N banshlaii* seems to be nearer on the basis of PC2. All the three species of *Nandus* form discrete group on PC1 vs. PC2.

**Fig. 3:**
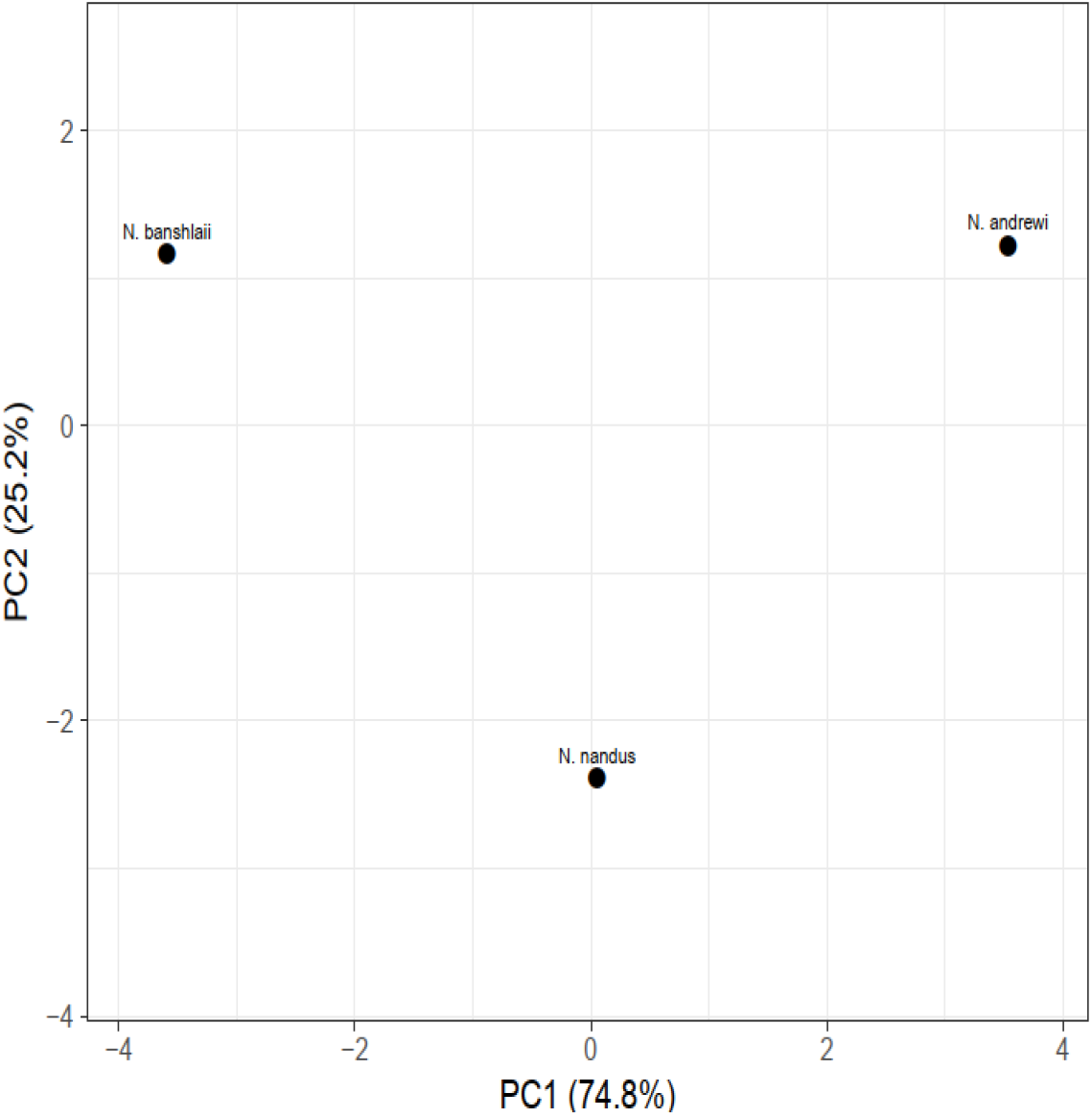
*N banshlaii* is separated from other two species of *Nandus* along first principal component, PC1 and PC2 contributed to 74.8% and 25.2% carnation respectively

**Table 3.** Comparative account on fin formula of seven species of *Nandus sp*.(Hossain and Sarker, 2013) and *N. banshlaii*.

| Parameters | *N. banshlaii* * | *N. meni* | *N. nandus* * | *N. oxyrhynchus* | *N. andrewi* * | *N. mercatus* | *N. nebulosus* | *N. prolixus* |
| --- | --- | --- | --- | --- | --- | --- | --- | --- |
| Dorsal fin | XII-XII,9-10 | XIII,11-13 | XII-XIV, 11-13 | XII,10 | XII-XIV, 11-14 | XIV,10 | XV-XVI, 10-12 | XIV,10 |
| Pectoral fin | 14-16 | 9-11 | I,15 | 14-15 | 14-16 | 15 | II,13-14 | 16 |
| Ventral fin | I,5 | I,7-13 | I,5 | I,6 | I,5-6 | I,5 | I,5 | I,5 |
| Anal fin | III,6-7 | III,7-18 | III,7-9 | III,5 | III,7-8 | III,5 - 6 | III,6-8 | III,6 |
| Caudal fin | 13-14 | 13-15 | 14 | 14 | 14-16 | 14 | 14 | 14 |
| Branchiostegal rays | 6 | 6 - 7 | 6 | 5 | 5 - 6 | 6 | 6 | 6 |
| References | Present study | Hossain & Sarker, 2013 | Talwar & Jhingran, 2001 | Ng et al.,1996 | Ng & Jaafar, 2008 | Ng, 2008 | Gray, 1835 | Chakrabarty et al., 2006 |

\*Indian species

*N. banshlaii* is distinctly separated from two other Indian species, *N. nandus* and *N. andrewi* on the basis of four morphological characters viz. predorsal length (PDL), prepectoral length (PPL), preventral length (PVL) and snout length (SN) while less distinctly by eye diameter (ED), interorbital distance (IOD), caudal peduncle depth in caudal peduncle length (CPDL) as shown in heatmap (Fig.4). Here *N. andrewi* is distinctly separated from other two species by caudal fin length (CFL), pre anal length (PAL) and head length (HL) and also less distinctly by dorsal fin base length (DFBL), anal fin base length (AFBL) and ventral fin length (VFL) but *N. nandus* is separated from other two species only on the basis of two characters, distinctly by body depth (BD) and indistinctly by caudal peduncle depth (CPD) (Fig. 3).

**Fig. 4:**
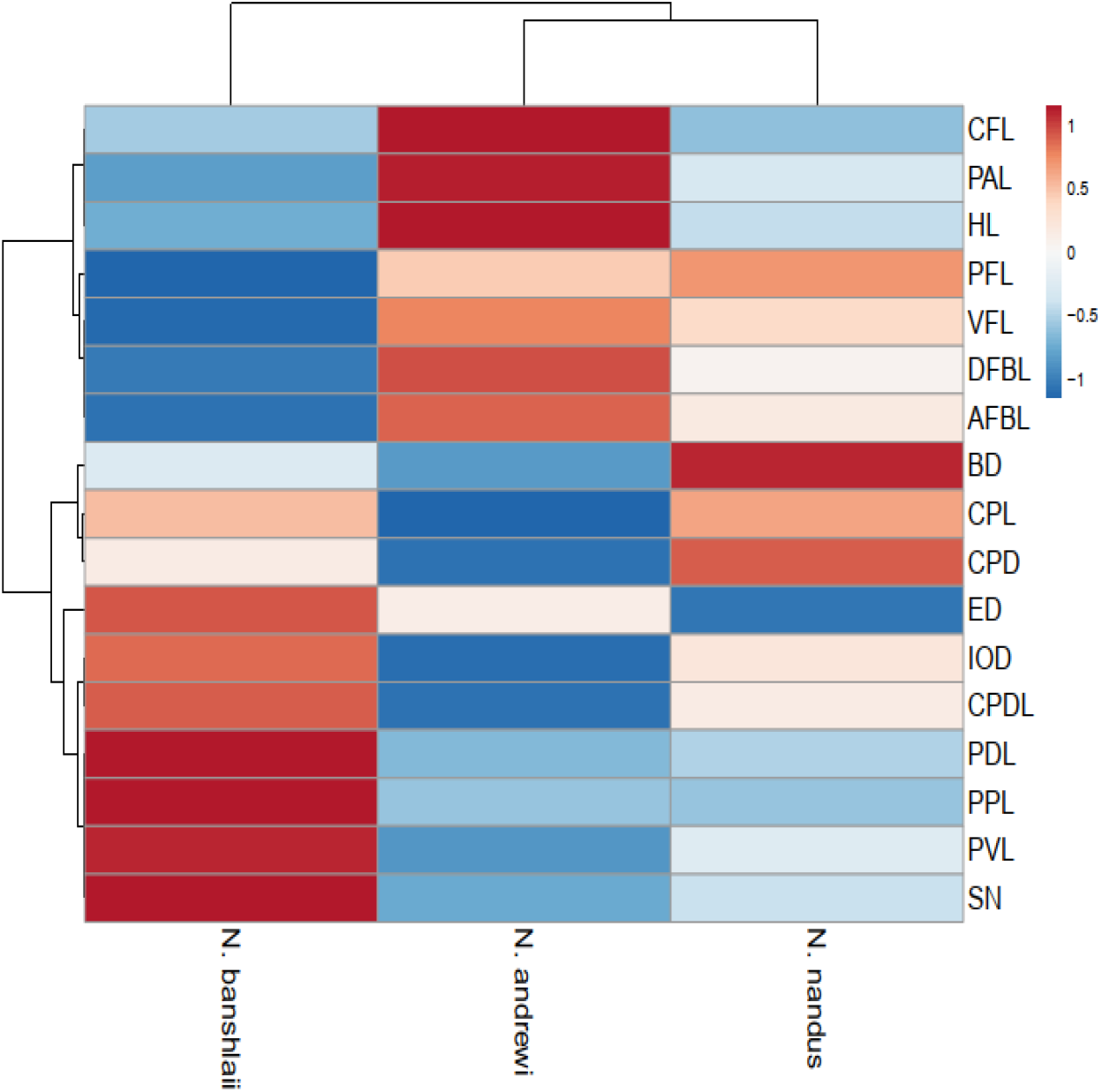
Heatmap generated using indices (along each row) for three different Indian species of *Nandus* using Clust Vis.

Dendrogram showed that *N. nandus* and *N. andrewi* are two closely related species and *N. banshlaii* is distantly related to both the species (Fig. 3).

## Discussion

Among eight species of *Nandus* so far reported, *N. nebulosus*, *N. oxyrhynchus*, *N. prolixus* and *N mercatus* distributed in different geographically isolated river basin, lack dark spot on caudal peduncle which is present in all the four species viz. *N. nandus, N andrewi, N. meni* and *N. banshlaii* of Ganga-Brahmaputra-Meghna river basin (of Gangetic delta). Similar observation is seen in the study of Tamang and Choudhry (2011) while describing *Glyptothorax dikrongensis* from Arunachal Pradesh. Among four species of *Glyptothorax*, distributed in two different river system (Ganga-Brahmaputra drainage and Salween drainage) characterized by presence or absence of banding pattern in the body, where they mentioned that morphological variations were due to different river basin those were geographically isolated. The present report supports this hypothesis.

Among all three Indian species, distribution of *Nandus nandus* in two separate river drainage i.e. Ganga-Brahmaputra and Irrawaddy, can be explained by palaeobiogeographic analyses of freshwater fishes. This analysis provides a link between the geological and biotic evolution of the Tibetan plateau and dispersion of fishes depends on the formation of direct connection between drainages (Bermingham and Martin, 1998; Lundberg, 1993). Almost similar distribution pattern is observed in *Gagata cenia* which is known from Ganga-Brahmaputra, Irrawaddy and Salween drainages (Chu et al., 1999; Karmakar, 2000; Roberts and Ferraris, 1998).

On the basis of morphometric characters, PCA and heatmap it can be concluded that *N. banshlaii* is distinctly separated and identified from all its congeners including Indian species also and should be treated as a new species of genus *Nandus*.

Since six years, we are vigorously surveying the ichthyofaunal diversity of Birbhum district (West Bengal, India) as this area is still to be explored properly. But there is no report of any species of *Nandus* from any water body in Birbhum except the present species. This reflects that *Nandus* species population is in precipitous decline at least in this area. Goswami and Dasgupta (2007) reported that *Nandus* population has declined to an alarming extent in West Bengal.

Key to three Indian species of *Nandus*:

1. Number of circumpeduncular scale 21 ---------- *N. andrewi*. More than 21 ---------- 2.
2. Shorter snout length (< 30% HL) ---------- *N. nandus*.
3. Longer snout length (> 30% HL) ---------- *N. banshlaii*.

## Comparative materials

### Nandus nandus

UMMZ 208417 (1), 60.9 mm SL; Bangladesh: Noakhali, Meghna River at island opposite Hajimara. UMMZ 208525 (1), 66.2 mm SL; Bangladesh: South Dakatia river at Faridganj, SE of Chandpur. UMMZ 208616 (1), 83.0 mm SL; Bangladesh: Comilla, Kunti Choumohani, P. S. Kaska, roadside ditch 27 Km south of Brahmabaria. UMMZ 208718 (4), 59.7-76.5 mm SL; Bangladesh: Comilla, Srma River at Lubachara, 51 Km ENE of Sylhet at Indian border. UMMZ 244771 (1), 58.2 mm SL; India: West Bengal, market at Barobisha. ZRC 39245 (1), 90.4 mm SL; India: West Bengal, Pulta. ZRC 50640 (2), 115.0 – 116.6 mm SL; India: West Bengal, Calcutta, Park Circus market. ZRC 51105 (2), 68.8 – 74.6 mm SL; India: West Bengal, Ichamati River drainage at Bangaon. ZRC 51107 (2), 43.9 – 47.4 mm SL; India: West Bengal, Ichamati River drainage in vicinity of Duttaphulia.

### Nandus andrewi

UMMZ 247483 (2), 77.3 – 78.0 mmSL, India: West Bengal. ZRC 51108 (1), 56.0 mm SL, India: West Bengal, Nadia district, Ichamati River drainage in the vicinity of Duttaphulia (23o14 N 88043 E).

### Nandus meni

MMSF 2013 E1 (Holotype) 11.9 mm SL, MMSF 2013 E2 113.1 mm SL, E3 119.6 mm SL, E4 100.8 mm SL, E5 117.3 mm SL, E6 94.1 mm SL, E7 112.0 mm SL, E8 113.1 mm SL, E9 109.6 mm SL, E10 100.8 mm SL, E11 117.3 mm SL, E12 94.2 mm SL, E13 111.9 mm SL, E14 113.1 mm SL, E15 109.6 mm SL, E16 111.9 mm SL. Bangladesh: Noakhali, freshwater swamp of Begumgonj (22^0^55’ N, 90^0^58’ E).

### Nandusprolixus

FMNH 44907 (holotype), 43.4 mm SL; FMNH 117232 (2 paratypes), 71.9 – 76.8 mm SL, Borneo: Sabah, Sandakan district, 26 km, north road Sandakan. FMNH 51964 (6 paratypes), 47.0 – 81.3 mm SL, Borneo: Sabah, 26 km north-west of Sandakan.

### Nandus mercatus

MZB 10987 (holotype), 70.0 mm SL, Sumatra, Sumatera Selatan, market at Sekayu, 2^0^51’ S 103^0^51’ E. ZRC 51419 (paratype), 39.8 mm SL, Sumatra: SumateraSelatan, S tributary of an Oxbow on the Musi River, in the vicinity of Danau Calak, 2057.190 S 103058.690 E.

### Nandus oxyrhynchus

ZRC 39246 (holoyype), 54.6 mm SL; ZRC 39247 (2 paratypes), 36.4 – 51.0 mm SL, Thailand: Sisaket, Amphoe Phrai Bung.UMMZ 218861 (21), 21.9 – 72.3 mm SL, Thailand: Khon Kaen, Nam Pong (Ubol Ratana) Reservoir, 2.5 km south of fish landing on the eastern shore. UMMZ 195557 (4), 66.8 – 72.0 mm SL, Thailand: Sakol Nakorn Fish Market. UMMZ 236920 (6), 55.0 – 70.7 mm SL, Thailand: Sun Chieng Mai in Kuran Payas.

### Nandus nebulosus

FMNH 16053 (1), 39.5 mm SL, Sumatra: Sumatera Selatan, Ogan River. UMMZ 246606 (1), 23.6 mm SL, Cambodia: Kampot, Prek Toek Sap, just above water falls; coastal drainage to the Gulf of Thailand. ZRC 33043 (1), 67.0 mm SL, Riau Archipelago; Pulau Bintan north. ZRC 37922 (2), 27.0 – 52.8 mm SL, Borneo: Sarawak, ditch 34 km from Mukah, on Mukah-Sibu road. ZRC 38570 (1), 36.1 mm SL, Sumatra: Jambi, Sungai Alai at 28 km on Muara Bango-Muara Tebo road. ZRC 39044 (1), 66.6 mm SL, Sumatra: Riau, Sungei Bengkwan. ZRC 39368 (1), 63.8 mm SL, Borneo: Sarawak, Sungei Stom Muda. ZRC 40833 (1), 61.0 mm SL, Thailand; Chantaburi, Klong Pheet. ZRC 42424 (1), 40.3 mm SL, Sumatra; Jambi, second stream on the road towards Muara Bulain after the junction or crossroads towards Palembang, Muara Bulian and Jambi. ZRC 42879 (4), 25.5 – 70.3 mm SL, Malaysia: Johar, Mersing. ZRC 51420 (4), 32.5 – 58.7 mm SL, Sumatra: Sumtera Selatan, Lalang drainage, swamp forest at Sentang, ca. 5 km after turnoff ca. 12 km after Bayung Lencir on Bayung Lencir-Jambi road, 1^0^56’ 11.5” S 103^0^42’ 31.9” E.

## Acknowledgement

Authors are likely to express their gratitude to the Teacher-in-Charge, Rampurhat College for his cooperation, Mr. Sangram Halder, Librarian, for technical assistance and Mr Soumendranath Bhakat of Lund University, Sweden for technical support (for PCA and heatmap analysis). We are thankful to Mr. Pradip Mandal and three students of our department, Master Supriyo Biswas, Master Sohel Aktar and Master Sandip Rajak for photo session.

---

**Plate 1.**
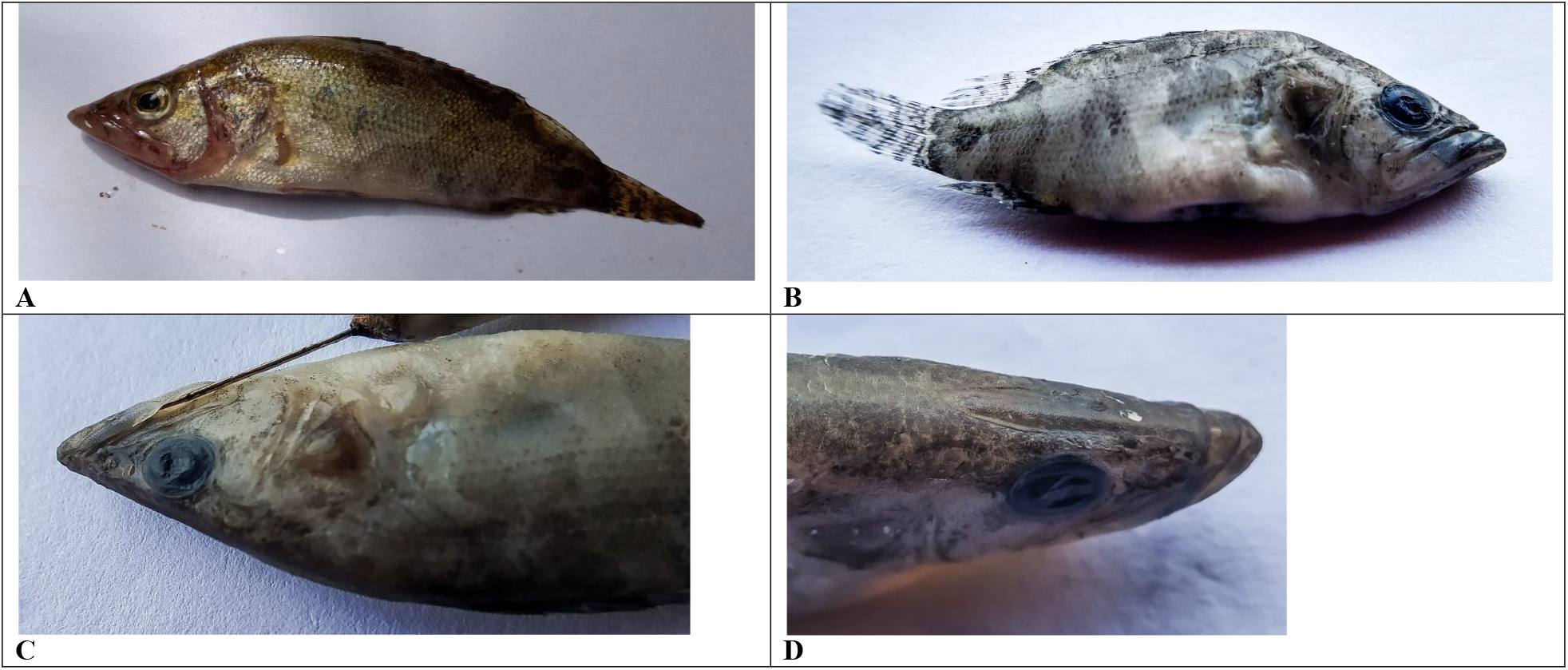
A. Live specimen of *Nandus banshlaii*. B. Preserved specimen showing Lateral line scale. C. Lower jaw showing semi lunar flap. D. Dorsal view of head showing long rod like ridge and projected lower jaw.

